# Aged-associated changes in active YAP staining during impaired skin wound healing in aged mice and delayed wound closure in ICE mice

**DOI:** 10.64898/2026.09.20.750311

**Authors:** Masayuki Tokunaga, Nao Kitamura, Mai Hamamoto, Motoshi Hayano, Kazuhiro Kimura, Makoto Furutani-Seiki, Yoichi Asaoka

**Author notes:** Masayuki Tokunaga and Nao Kitamura contributed equally to this work.

## Abstract

Aging is associated with impaired skin wound healing, but the molecular mechanisms underlying this decline remain incompletely understood. In particular, how aging affects the temporal regulation of mechanosensitive signaling during tissue repair is poorly understood. In this study, we investigated age-associated changes in cutaneous wound healing, with a focus on Yes-associated protein (YAP)-associated mechanosensitive responses, and examined whether ICE mice, an inducible model of epigenetic aging, reproduce age-associated impairment in wound repair. Splinted full-thickness excisional wounds were generated in young and naturally aged mice, and wound closure, gene expression, and active YAP were evaluated over time. Aged mice exhibited delayed wound closure, particularly during the mid-to-late phases of repair. Expression of the fibrosis-related genes *Tgfb1*, *Col1a1*, and *Col3a1*, and the YAP-associated genes *Ctgf*, *Cyr61*, and *Arhgap18* did not differ significantly between young and aged wounds at the examined time points. In contrast, *Ankrd1* was selectively elevated in aged wounds at postoperative day 14. Active YAP immunofluorescence showed age-associated differences in staining patterns during wound healing, indicating altered regulation of mechanosensitive signaling with aging. Furthermore, ICE mice exhibited delayed wound closure compared with Cre control mice, reproducing a major functional feature observed in naturally aged mice. These findings suggest that age-associated impairment of wound healing is characterized not by uniform activation or suppression of YAP-related signaling, but by altered temporal regulation of mechanosensitive responses together with selective *Ankrd1* elevation. ICE mice could provide a useful experimental model for investigating age-associated defects in cutaneous wound healing.

## Introduction

Aging is characterized by a progressive decline in tissue and organ function, leading to increased vulnerability to disease and reduced health span (López-Otín et al. 2013). At the cellular and molecular levels, aging is associated with multiple interconnected processes, including genomic instability, epigenetic alterations, mitochondrial dysfunction, cellular senescence, chronic inflammation, and altered intercellular communication, which have been conceptualized as hallmarks of aging (López-Otín et al. 2023). Among the functional consequences of aging, impaired skin wound healing is clinically important (Tobin 2017; Khalid et al. 2022). Aging delays multiple steps of wound repair, including inflammatory resolution, re-epithelialization, granulation tissue formation, and extracellular matrix remodeling (Landén et al. 2016; Peña and Martin 2024; Mamun et al. 2024). However, the molecular mechanisms that link aging to delayed wound repair remain poorly understood, particularly with respect to how signaling activities are dynamically regulated over time during the healing process.

Yes-associated protein (YAP) and transcriptional coactivator with PDZ-binding motif (TAZ) are transcriptional coactivators regulated by the Hippo pathway that are key mediators of mechanotransduction and regulate diverse biological processes, including cell migration, proliferation, tissue growth, and three-dimensional tissue morphogenesis (Dupont et al. 2011; Porazinski et al. 2015; Moya and Halder 2019). In skin wound healing, YAP/TAZ activity has been implicated in tissue repair, as knockdown of YAP and TAZ delays wound closure in mice, supporting a requirement for YAP/TAZ activity during skin repair (Lee et al. 2014). In addition, Yap1 promotes epidermal stem cell proliferation and tissue expansion downstream of α-catenin (Schlegelmilch et al. 2011). However, YAP activation does not appear to have a uniformly beneficial effect on wound repair. Although pharmacological YAP activation using PY-60 has been reported to promote regenerative repair of skin wounds in pig and human skin models, excessive or sustained YAP activation may promote abnormal tissue growth or hyperproliferative responses (Grzelak et al. 2023). Conversely, inhibition of mechanotransduction through YAP blockade in wound fibroblasts has been reported to reduce scarring and promote regenerative repair (Mascharak et al. 2021). These findings suggest that YAP activity during wound healing is not simply beneficial or detrimental, but is likely regulated in a context-dependent and temporally dynamic manner. Therefore, understanding the temporal dynamics of active YAP during wound healing may be important for distinguishing productive tissue repair from impaired or fibrotic repair.

ANKRD1 (ankyrin repeat domain 1) is a stress- and mechanosensitive transcriptional regulator that has been implicated in tissue injury and repair. In skin, ANKRD1 expression is strongly induced after wounding in multiple cell types, including keratinocytes, endothelial cells, inflammatory cells, and fibroblasts (Shi et al. 2005). Functional studies further showed that loss of Ankrd1 delays wound closure and impairs granulation tissue formation, fibroblast migration, and collagen matrix contraction, supporting a role for ANKRD1 in coordinating cellular responses during tissue repair (Samaras et al. 2015). More recently, ankrd1a was shown to be persistently upregulated in cardiomyocytes bordering the injury or scar during zebrafish heart regeneration and to modulate their dedifferentiation, further suggesting a role for ANKRD1 in injury-associated cellular remodeling (Boskovic et al. 2026). Together, these findings identify ANKRD1 as an injury-responsive factor associated with tissue remodeling and regeneration.

Although comparisons between young and naturally aged mice are useful for identifying age-associated changes in wound repair, such comparisons can be influenced by differences in lifespan, housing duration, systemic physiology, and environmental exposure (Flurkey et al. 2007). Therefore, complementary models are needed to examine how aging-associated molecular alterations affect tissue repair. The inducible changes to the epigenome (ICE) mice have been developed as an inducible model of epigenetic aging, in which transient DNA double-strand breaks induce epigenetic alterations during DNA repair and thereby accelerate aging-like phenotypes (Yang et al. 2023). This model provides an opportunity to investigate how DSB-induced epigenetic aging affects the regulation of mechanosensitive signaling during tissue repair. Here, we investigated whether aging alters the temporal dynamics of YAP activity during skin wound healing. We also used ICE mice as a complementary model to assess wound healing in the context of aging-like phenotypes induced by DNA double strand breaks.

## Results

### Aged mice exhibit delayed cutaneous wound closure

To examine the effect of aging on skin wound healing, we generated splinted full-thickness excisional wounds on the dorsal skin of young and aged mice. Wound closure increased progressively in both groups, but aged mice exhibited a slower closure rate compared with young mice. Wound closure was significantly delayed in aged mice, with aged wounds remaining less closed than young wounds from postoperative day (POD) 3 to POD14 (Fig. 1). These findings demonstrate that aged mice show delayed skin wound closure.

**Fig. 1.**
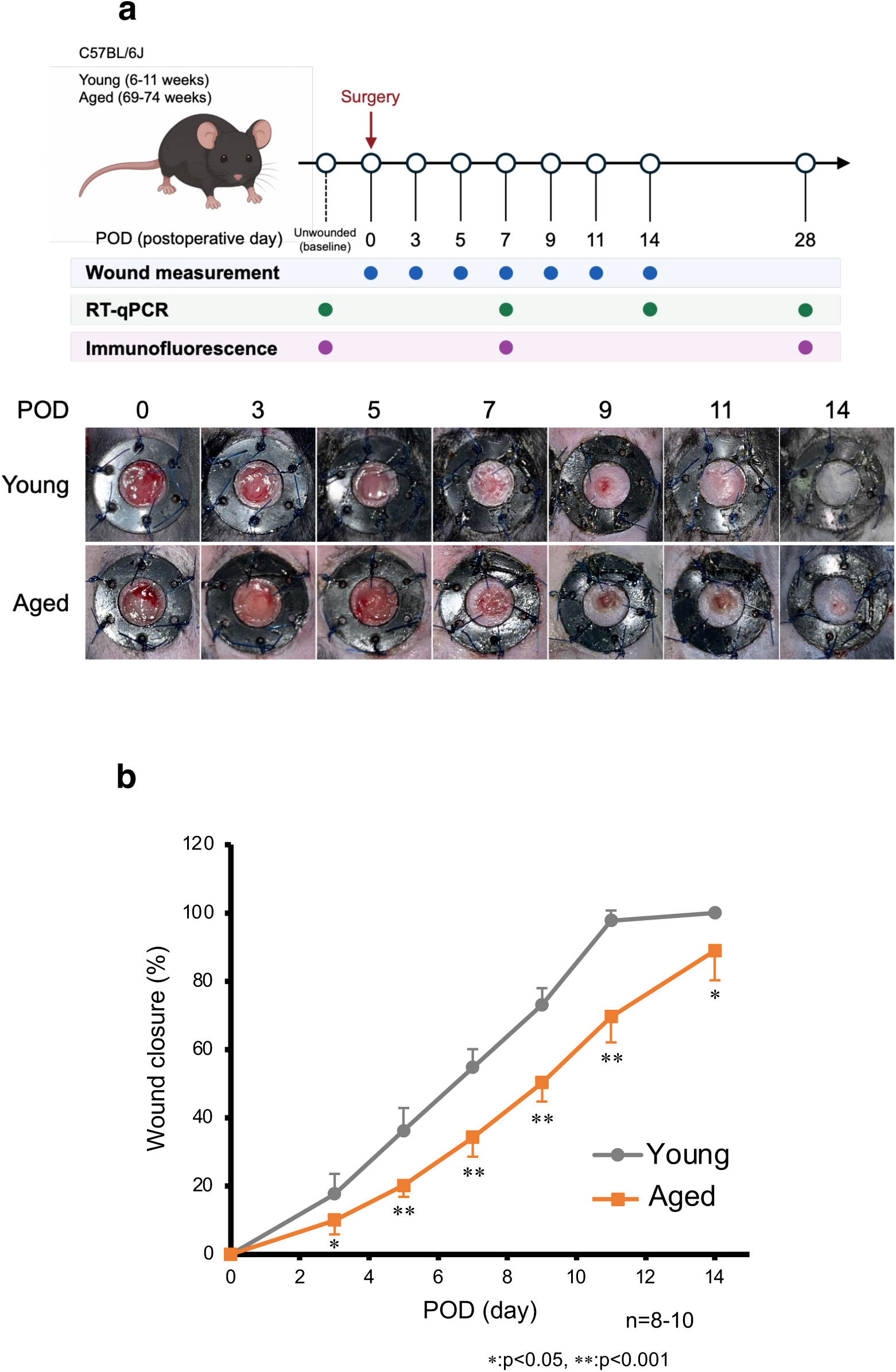
Aged mice show delayed skin wound closure. **a** Experimental scheme of the wound healing assay in young and aged mice. Representative macroscopic images of wounds from young and aged mice at the indicated postoperative days (PODs) are shown. **b** Quantification of wound closure over time. Aged mice showed significantly delayed wound closure compared with young mice from POD3 to POD14. Data are shown as mean ± SD; n = 8–10 wounds per group. Statistical significance was determined by Tukey’s HSD test for least-squares mean differences. *P < 0.05. **P < 0.001.

### Aged wounds show selective *Ankrd1* elevation and age-associated differences in active YAP staining

To explore molecular changes associated with delayed wound closure in aged mice, we analyzed the expression of fibrosis-related and YAP/mechano-homeostasis-related genes by RT-qPCR. Among the fibrosis-related genes examined, including *Tgfb1*, *Col1a1*, and *Col3a1*, no significant differences were detected between young and aged wounds at the analyzed time points. Similarly, the expression of *Ctgf*, *Cyr61*, and *Arhgap18* did not differ significantly between the two groups (Fig. 2).

**Fig 2.**
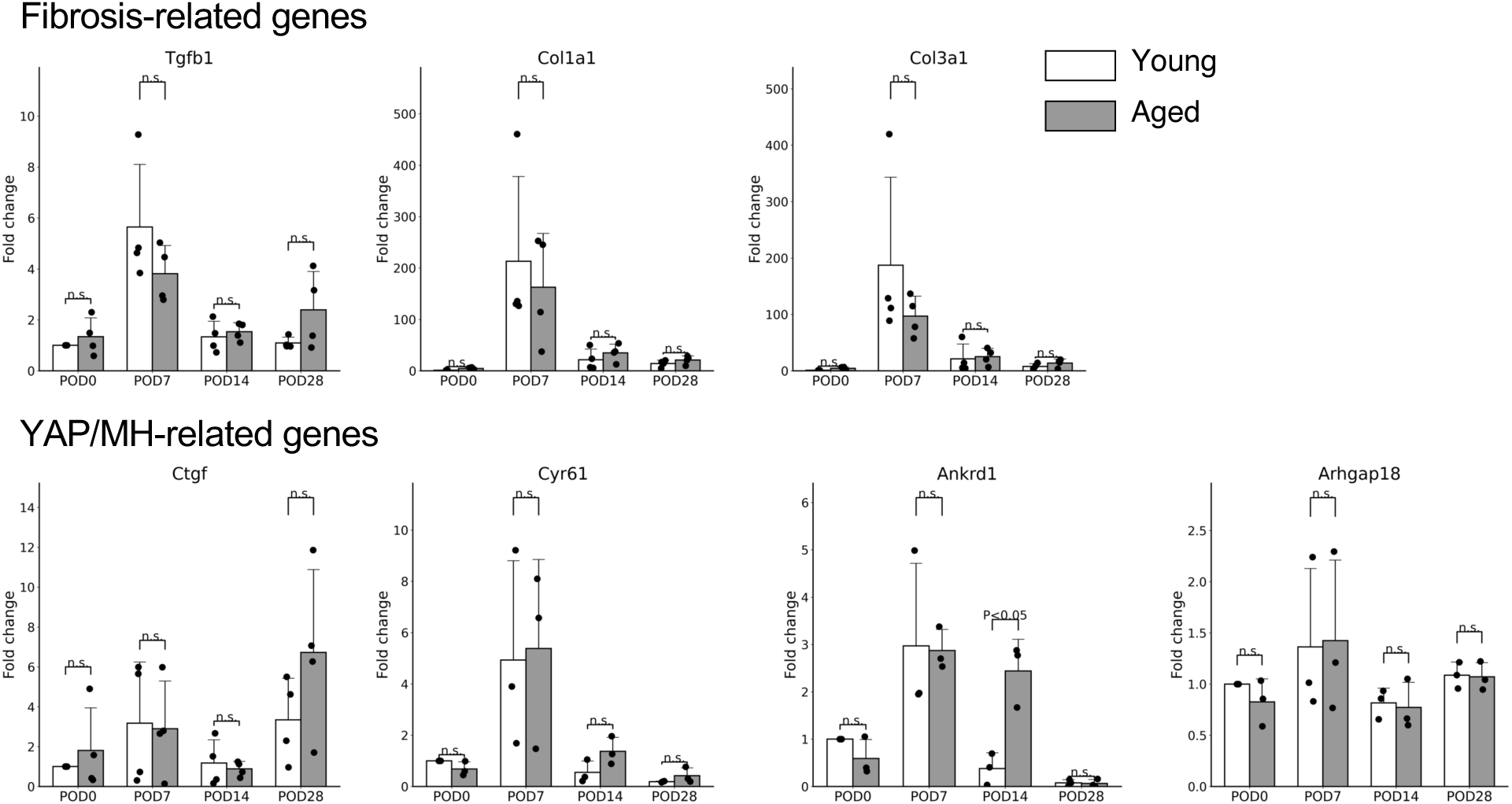
*Ankrd1* is selectively elevated in aged wounds at POD14. RT-qPCR analysis of fibrosis-related genes and YAP/mechano-homeostasis-related genes in young and aged wounds at the indicated postoperative days. The expression levels of *Tgfb1*, *Col1a1*, *Col3a1*, *Ctgf*, *Cyr61*, and *Arhgap18* were not significantly different between young and aged wounds at the analyzed time points. In contrast, *Ankrd1* expression was significantly increased in aged wounds compared with young wounds at POD14. Data are shown as mean ± SD; n = 4 mice per group. Statistical significance was determined by Tukey’s HSD test for least-squares mean differences. *P < 0.05; n.s., not significant.

In contrast, *Ankrd1* showed a selective age-dependent alteration. *Ankrd1* expression was significantly increased in wounds of aged mice compared to young mice at POD14, whereas no significant difference was observed at POD0 (unwounded), POD7, or POD28. These results indicate that wounds of aged mice do not exhibit upregulation of fibrosis-related genes or canonical YAP-associated target genes. Rather, *Ankrd1* was selectively elevated at POD14, indicating that *Ankrd1* expression is maintained or enhanced during mid-to-late phases of wound repair in aged mice (Fig. 2). Because *Ankrd1* is a YAP-responsive gene, we next examined whether YAP activation itself was also altered during wound healing.

To directly assess the spatial and temporal distribution of YAP activation during wound healing, we performed immunofluorescence staining using an antibody recognizing active YAP. In young mice, active YAP-positive cells were abundant in unwounded skin at POD0 (unwounded) with less prominent staining at later stages of wound healing. In contrast, aged mice showed very few active YAP-positive cells at POD0 (unwounded), whereas more prominent staining was observed at later time points, including POD7 and POD28. These observations suggested aged-associated differences in the temporal pattern of active YAP staining during wound healing (Fig. 3).

**Fig. 3.**
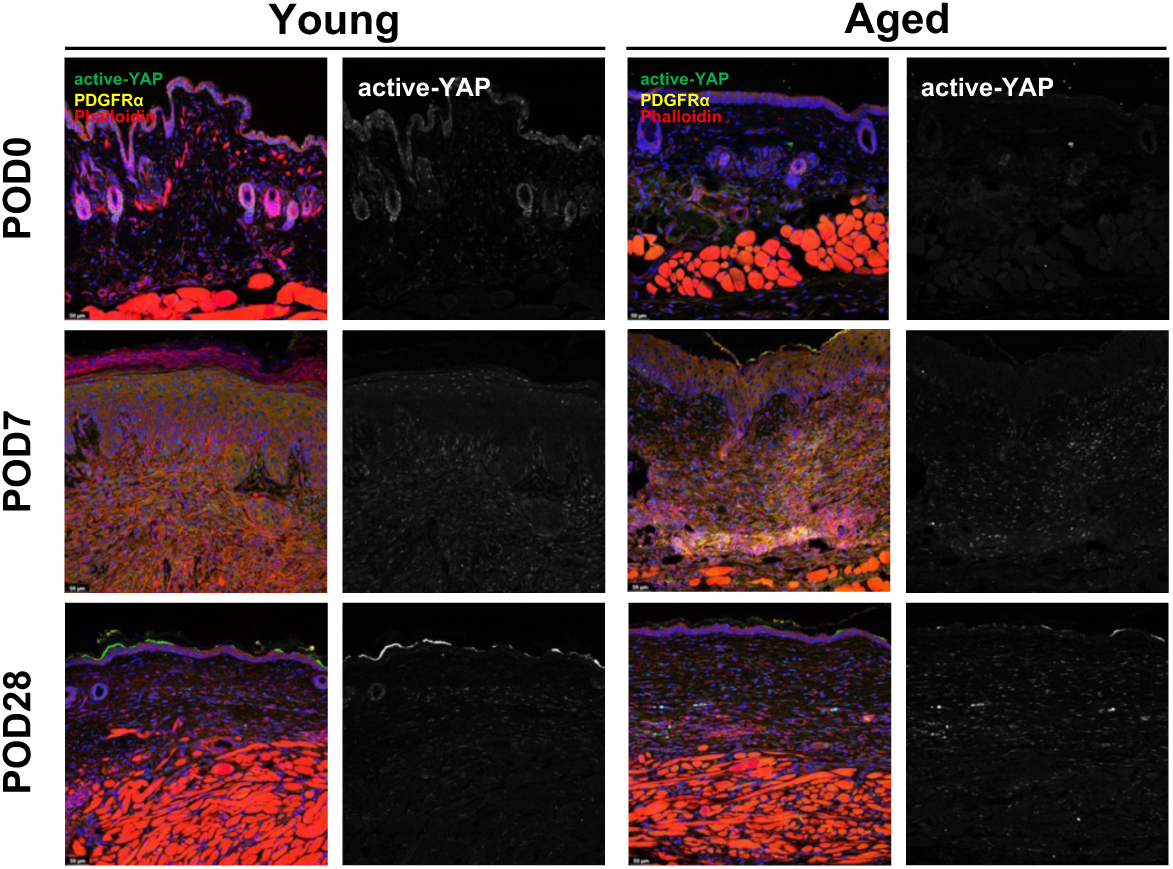
Active YAP immunofluorescence in young and aged wounds. Representative immunofluorescence images of active YAP, PDGFRα, phalloidin, and nuclei in young and aged wounds at POD0 (unwounded), POD7, and POD28. Grayscale images show the active YAP channel alone. In young wounds, active YAP staining appeared lower at later stages, whereas aged wounds showed weak staining at POD0 (unwounded) and more prominent staining at later time points.

### ICE mice show delayed cutaneous wound closure

To test whether DSB-associated epigenetic aging recapitulates age-associated defects in wound repair, we subjected Cre control and ICE mice to splinted full-thicknessexcisional wounding. Cre control and ICE mice were fed a tamoxifen-containing diet for 4 weeks at 4–6 months of age to induce I-*Ppo*I expression. After a 7–10-month post-treatment period, wound closure was monitored over time. Compared with Cre control mice, ICE mice exhibited delayed wound closure, with significantly reduced closure rates from POD3 through POD14 (Fig. 4). These findings suggest that ICE mice display impaired skin wound repair, consistent with the delayed wound closure observed in naturally aged mice.

**Fig 4.**
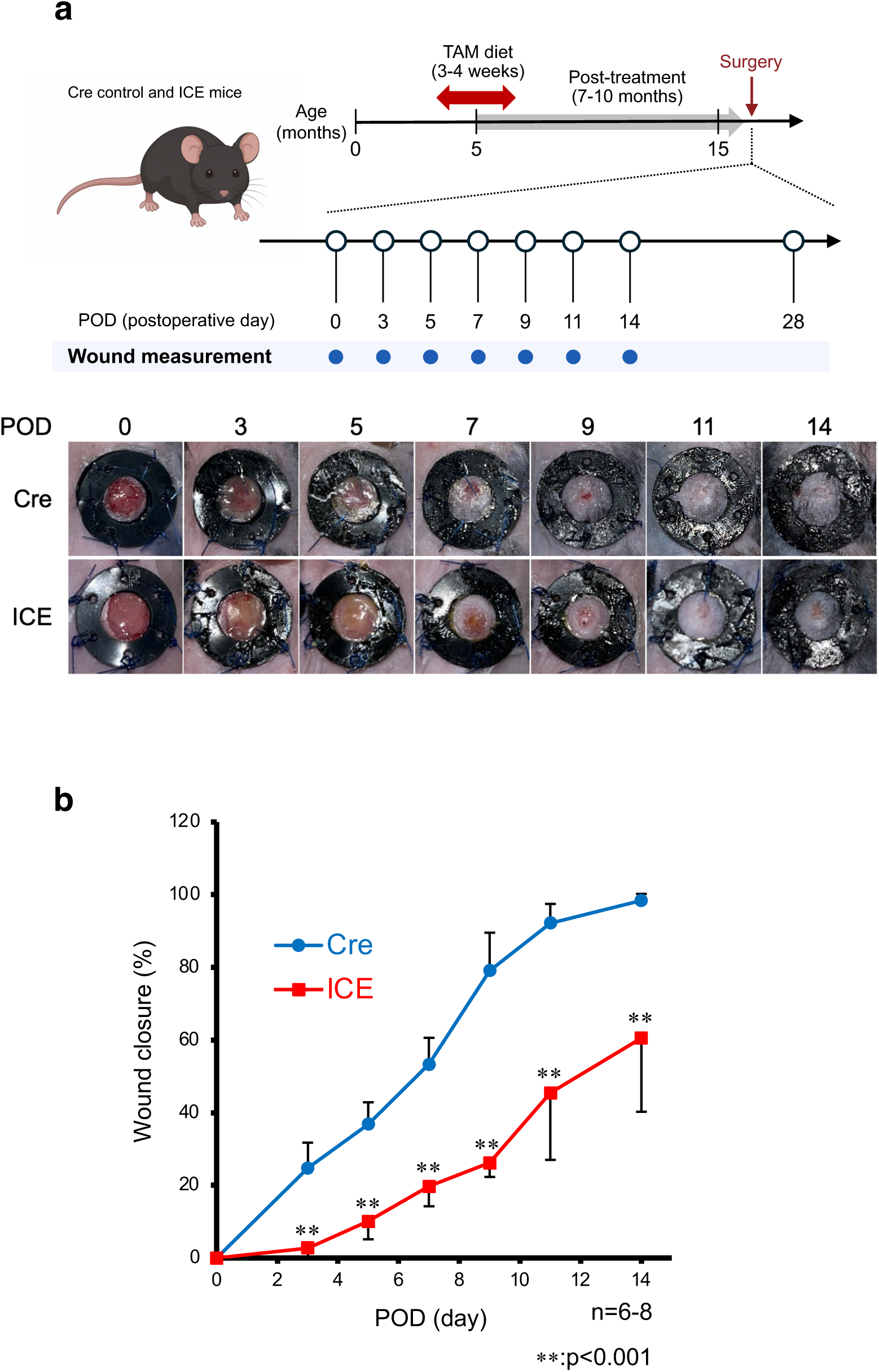
ICE mice show delayed skin wound closure. **a** Experimental scheme of the wound healing assay in Cre control and ICE mice. Representative macroscopic images of wounds from Cre control and ICE mice at the indicated PODs are shown. **b** Quantification of wound closure over time. ICE mice showed significantly delayed wound closure compared with Cre control mice. Data are shown as mean ± SD. n = 6–8 wounds per group. Statistical significance was determined by Tukey’s HSD test for least-squares mean differences. **P < 0.001.

## Discussion

In this study, we investigated how aging affects cutaneous wound healing, with particular focus on the temporal regulation of YAP-associated mechanosensitive signaling and evaluated whether ICE mice can serve as a useful model of age-associated impairment in wound repair. Naturally aged mice exhibited delayed wound closure compared with young mice, particularly during the mid-to-late phases of repair. This functional impairment was accompanied by a selective elevation of *Ankrd1* at POD14 and an age-dependent alteration in active YAP staining. In contrast, the fibrosis-related genes *Tgfb1*, *Col1a1*, and *Col3a1*, as well as the YAP-associated genes *Ctgf*, *Cyr61*, and *Arhgap18*, did not differ significantly between young and aged wounds at the examined time points. Furthermore, ICE mice, in which transient DNA double-strand breaks induce persistent epigenetic alterations and aging-like phenotypes, also exhibited delayed wound closure. Together, these findings suggest that impaired wound healing with aging is associated not with a uniform enhancement or suppression of YAP-related signaling, but rather with altered temporal regulation of mechanosensitive responses during tissue repair, and that ICE mice could be a useful experimental model for investigating age-associated defects in wound healing.

Several limitations should be considered. First, gene expression was analyzed in whole wound tissue, potentially masking cell type-specific changes in YAP-associated and fibrotic pathways. Second, the transient transcriptional responses between the analyzed time points may have been missed. Cell type-resolved analyses, such as co-localization with cell type-specific markers and single-cell transcriptomic profiling, will help define the cellular origins and functional consequences of the age-dependent changes observed here. Finally, although ICE mice recapitulated the delayed wound closure observed in naturally aged mice, further molecular characterization is needed to determine the extent to which the underlying mechanisms, including alterations in mechanosensitive signaling, are shared between the two models.

In conclusion, our findings demonstrate that aging is associated with delayed cutaneous wound closure and altered temporal regulation of YAP-associated mechanosensitive responses. Rather than exhibiting a generalized increase in fibrotic or canonical YAP target gene expression, aged wounds showed a selective increase in *Ankrd1* expression during the mid-to-late phase of repair, together with altered active YAPstaining. The similar delay in wound closure observed in ICE mice suggests that this model may provide a useful platform for investigating age-associated defects in wound healing.

### Data availability

All data related to this published article is available from the corresponding author and upon reasonable request.

## Materials and methods

### Animals and ICE mice induction

C57BL/6J and I-*Ppo*I^STOP/+^, Cre^ERT2/+^ mice were used in this study. Female young C57BL/6J mice (6-11-week-old) and aged C57BL/6J mice (69-74-week-old) were purchased from Japan SLC (Shizuoka, Japan). The I-*Ppo*I^STOP/+^ and Cre^ERT2/+^ mouse lines were originally established as previously described (Yang et al. 2023). ICE mice were generated by crossing I-*Ppo*I^STOP/+^ and Cre^ERT2/+^. Male 4-6-month-old Cre control and ICE mice were fed a modified AIN-93G purified rodent diet containing 360 mg/kg tamoxifen citrate (Research Diets, D21042909) for 3 or 4 weeks to induce I-*Ppo*I expression. Wound healing assays were performed 7–10 months after completion of tamoxifen treatment. All mice were maintained at the Science Research Center, Institute of Life Science and Medicine, Yamaguchi University at constant temperature, humidity and light/dark cycle. Mice had ad libitum access to food and water, except during administration of tamoxifen-containing diet.

### Skin wound healing assay

Skin wound healing assays were performed under anesthesia. Mice received buprenorphine (0.2 mg/kg, s.c., Nissin Pharmaceutical) before surgery for analgesia. General anesthesia was induced by intraperitoneal administration of a three-component anesthetic mixture consisting of medetomidine (0.3 mg/kg, Domitor, Nippon Zenyaku Kogyo), midazolam (4 mg/kg, Maruishi Pharmaceutical), and butorphanol (5 mg/kg, Vetorphale, Meiji Animal Health), administered at 0.1 mL per 10 g body weight. After induction of anesthesia, dorsal hair was removed using an electric clipper and depilatory cream, and the dorsal skin was disinfected with 70% ethanol. Two full-thickness excisional wounds were generated on the dorsal skin using a 4-mm biopsy punch (Kai Industries, BPP-40F). Wounds were positioned bilaterally at the same rostrocaudal level, at least 6 mm caudal to the base of the ears and approximately 4 mm lateral to the dorsal midline. After marking the wound area with the biopsy punch, the skin was excised using surgical scissors. To minimize wound contraction, a silicone wound splint (CSCRIE, R-CSC086-01) was attached around each wound using Vetbond (3M, 10219512) tissue adhesive and secured with eight interrupted sutures using 6-0 Ethilon (alfresa pharma, HR1605NA75-KF2). The wound surface was covered with a Tegaderm Transparent dressing (3M, 8-1896-01). Dressings were replaced every other day under inhalation anesthesia until postoperative day 11. Buprenorphine (0.2 mg/kg, s.c.) was administered again 4–6 h after surgery and twice on the following day.

### Wound closure analysis

Wounds were photographed under inhalation anesthesia on PODs 3, 5, 7, 9, 11, and 14. Images were acquired with a ruler placed in the same field of view for calibration. At each time point, the long and short diameters of each wound were measured using calipers.

Wound area was estimated using the formula for an ellipse: area = π × long diameter / 2 × short diameter / 2. Wound closure was expressed as the percentage reduction in wound area relative to the initial wound area measured immediately after injuring, using the formula: wound closure (%) = [(initial wound area – wound area at each time point) / initial wound area] x 100.

### Tissue collection and sample preparation

Wound tissues were collected from wound sites on PODs 7, 14, and 28 and from unwounded control skin (POD0). For samples collected on POD14 and POD28, regenerated hair around the wound area was removed using an electric clipper before tissue collection. The wound area was excised using an 8-mm skin biopsy punch (Kai industries, BP-80F) and divided into two halves through the center of the wound. The one half of each sample was placed in a 1.5-mL tube, immediately frozen in liquid nitrogen, and stored at −80°C until RNA extraction. The other half was immersed in 4% paraformaldehyde and fixed at room temperature for immunofluorescence analysis. For unwounded control skin (POD0) samples, uninjured dorsal skin was collected in the same manner.

After fixation for 48 h, tissues were washed three times with PBS for 5 min each. Samples were then cryoprotected by sequential immersion in 10%, 20%, and 30% sucrose in PBS at 4°C with gentle agitation until the tissues sank. After cryoprotection in 30% sucrose, tissues were embedded in a mixture of O.C.T. Compound (Sakura Finetek) and 20% sucrose at a 2:1 ratio (v/v) and stored at −80°C until cryosectioning.

### RNA extraction and RT-qPCR

Total RNA was extracted from wound tissues using TRIzol reagent (Invitrogen, 15596018). Briefly, frozen skin samples were minced into small pieces and homogenized in 1 mL TRIzol reagent using a μT-12 bead crusher (Taitec) with one 5-mm stainless steel bead (TAITEC, 0068220-000) in a 2.0-mL tube (Ina-optika, 2641-0B). Homogenization was performed for three cycles of 30 s at 3,200 rpm, with cooling on ice for 1 min between cycles. Samples were incubated at room temperature for 5 min, followed by phase separation with 200 μL chloroform (Sigma, 05-3400-5). After centrifugation at 12,000 × g for 15 min, the aqueous phase was collected and subjected to a second chloroform extraction. RNA was precipitated by adding 1 μL glycogen solution (20 mg/mL, Nacalai, 17110-11) and an equal volume of isopropanol (FUJIFILM Wako, 168-21675), followed by incubation at room temperature for 10 min. After centrifugation at 12,000 × g for 10 min, RNA pellets were washed twice with 70% ethanol (FUJIFILM Wako, 059-07895), air-dried briefly, and dissolved in RNase-free water. RNA samples were incubated at 55°C for 10 min to ensure complete dissolution and stored at −80°C until use. RNA concentration and purity were assessed using a NanoDrop One spectrophotometer (Thermo Fisher Scientific).

Complementary DNA was synthesized from 500 ng of total RNA using ReverTra Ace qPCR RT Master Mix with gDNA Remover (TOYOBO, FSQ-301) according to the manufacturer’s instructions. Quantitative PCR was performed using PowerUp SYBR Green Master Mix for qPCR (Thermo Fisher Scientific, A25742) or SYBR Green qPCR Master Mix (Thermo Fisher Scientific, A66732) on a QuantStudio 3 Real-Time PCR System (Thermo Fisher Scientific). The expression levels of *Tgfb1*, *Col1a1*, *Col3a1*, *Ctgf*, *Cyr61*, *Ankrd1*, and *Arhgap18* were analyzed. For normalization, the mean Ct value of *Rpl13a* and *Rplp0* was used as the reference Ct. Relative gene expression was calculated using the ΔΔCt method. Primer sequences are listed in Supplementary Table1.

### Immunofluorescence staining and Image acquisition

Frozen wound tissues embedded in O.C.T. Compound were sectioned at 10-μm thickness using a cryostat (Leica, CM1950) and mounted on glass slides (Matsunami glass, SMAS-01). Cryosections were brought to room temperature and washed with PBS to remove residual O.C.T. Compound. Sections were permeabilized and blocked in PBS containing 0.2% Triton X-100 and 5% normal goat serum (Abcam, ab7481) for 30 min at room temperature. Sections were then incubated overnight at 4°C with primary antibodies against active YAP (rabbit, Abcam, ab205270, 1:100) and CD140a/PDGFRα (rat, Invitrogen, 14-1401-81, 1:100) diluted in blocking buffer.

After washing with 0.05% PBST, sections were incubated for 2 h at room temperature in the dark with Alexa Fluor 488 goat anti-rabbit IgG (Thermo Fisher Scientific, A-11034, 1:250) and Alexa Fluor 555 goat anti-rat IgG (Thermo Fisher Scientific, A-21434, 1:250). F-actin was stained with Phalloidin-iFluor 647 reagent (Abcam, ab176759, 1:1000), and nuclei were counterstained with DAPI solution (Dojindo, D523, 1:1000). Sections were mounted with SlowFade Diamond Antifade Mountant (Invitrogen, S36963) and coverslipped.

Fluorescence images were acquired using a STELLARIS 8 confocal microscope (Leica Microsystems) equipped with 5× or 20× objectives. Images were captured from the wound bed region under identical acquisition settings within each staining batch. For quantitative analysis, three fields per wound were acquired from each mouse. All images used for comparison were acquired using identical laser power, gain, offset, and other acquisition settings.

For active YAP immunofluorescence, samples were stained in balanced batches. Each staining batch included samples from all time points and both age groups to minimize confounding between staining batch and postoperative day.

### Statistical analysis

Statistical analyses for wound closure and RT-qPCR data were performed using JMP Student (version 19.1.0, SAS Institute Inc., Cary, NC, USA). Data are presented as mean ± SD. For wound closure analysis, a mixed-effects model was used to assess the effects of group, postoperative day, and their interaction, followed by Tukey’s HSD test for least-squares mean differences between groups at each postoperative day. For RT-qPCR analyses, a two-way linear model was used to assess the effects of group, postoperative day, and their interaction, followed by Tukey’s HSD test for least-squares mean differences. A value of P < 0.05 was considered statistically significant.

## Acknowledgments

We thank the members of the Department of Systems Biochemistry in Pathology and Regeneration, Yamaguchi University Graduate School of Medicine, for valuable help. We thank Science Research Center, Institute for Biomedical Research and Education for their facilities and support, including the use of STELLARIS 8 confocal microscope, Cryostat and QuantStudio3. During the preparation of this manuscript, the authors used ChatGPT (OpenAI) to assist with language editing and manuscript organization. All scientific interpretations and conclusions were reviewed and determined by the authors, who take full responsibility for the content of the publication.

## Funding

This work was supported by a Grant from the Japanese Ministry of Education, Culture, Sports, Science, and Technology, 24K02226 for M.F.S.

## Author contributions

Y. Asaoka and M. Furutani-Seiki conceived the project. M. Tokunaga, N. Kitamura, M. Furutani-Seiki, and Y. Asaoka contributed to the design of the experiments. M. Tokunaga, N. Kitamura and M. Hamamoto performed the experiments and contributed to the collection and analysis of data. N. Kitamura conducted the statistical analysis. M. Hayano and K. Kimura contributed to the interpretation of the results and scientific discussions. M. Tokunaga, N. Kitamura, M. Hayano, M. Furutani-Seiki, and Y. Asaoka contributed to the writing of the manuscript. All authors reviewed and approved the final manuscript.

## Ethical Approval

All animal procedures were conducted following the guidelines approved by the Committee for the Ethics of Animal Experiments at the Yamaguchi University School of Medicine.

## Competing interests

The authors declare no competing interests.

## Supplementary Figure legends

**Supplementary Table 1.**
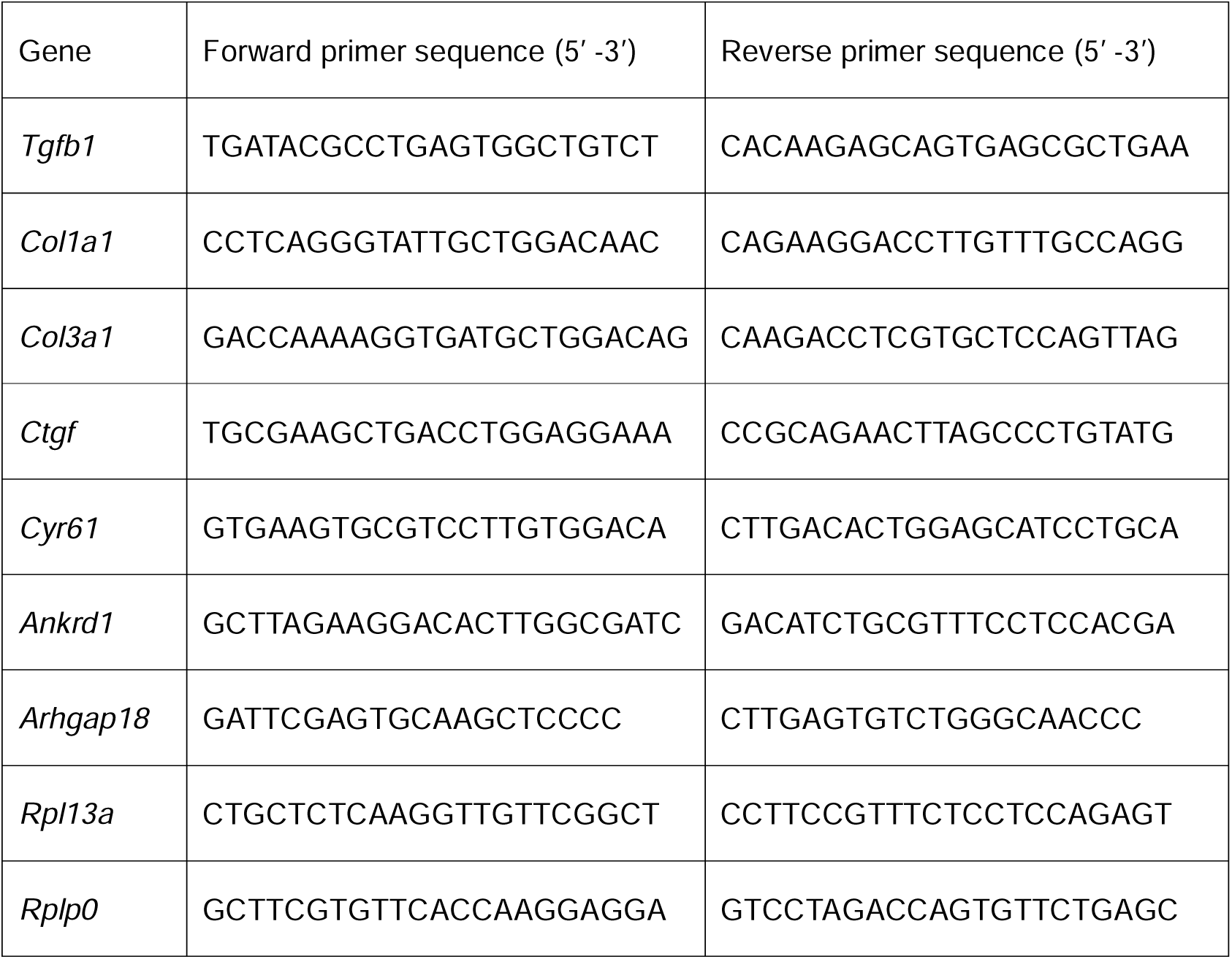
Primer sequences used for RT-qPCR.

## Notes

### Competing Interest Statement

The authors have declared no competing interest.

## References

Boskovic S, Novkovic M, Milicevic A, Milosevic E, Jasnic J, Milovanovic M, Kojic S (2026) ankrd1a is consistently upregulated in cardiomyocytes bordering the injury or scar area and affects their dedifferentiation during zebrafish heart regeneration. Cells & Development 186:204082. 10.1016/J.CDEV.2026.204082

Dupont S, Morsut L, Aragona M, Enzo E, Giulitti S, Cordenonsi M, Zanconato F, Le Digabel J, Forcato M, Bicciato S, Elvassore N, Piccolo S (2011) Role of YAP/TAZ in mechanotransduction. Nature 474:179–183. 10.1038/nature10137

Flurkey K, Currer JM, Harrison DE (2007) Mouse Models in Aging Research. The Mouse in Biomedical Research 3:637–672. 10.1016/B978-012369454-6/50074-1

Grzelak EM, Dayan Elshan NGR, Shao S, Bulos ML, Joseph SB, Chatterjee AK, Chen JJ, Nguyên-Trân V, Schultz PG, Bollong MJ (2023) Pharmacological YAP activation promotes regenerative repair of cutaneous wounds. Proc Natl Acad Sci U S A 120:e2305085120. 10.1073/PNAS.2305085120

Khalid KA, Nawi AFM, Zulkifli N, Barkat MA, Hadi H (2022) Aging and Wound Healing of the Skin: A Review of Clinical and Pathophysiological Hallmarks. 12:2142. 10.3390/LIFE12122142

Landén NX, Li D, Ståhle M (2016) Transition from inflammation to proliferation: a critical step during wound healing. Cellular and Molecular Life Sciences 73:3861–3885. 10.1007/S00018-016-2268-0

Lee MJ, Byun MR, Furutani-Seiki M, Hong JH, Jung HS (2014) YAP and TAZ Regulate Skin Wound Healing. Journal of Investigative Dermatology 134:518–525. 10.1038/JID.2013.339

López-Otín C, Blasco MA, Partridge L, Serrano M, Kroemer G (2013) The Hallmarks of Aging. Cell 153:1194–1217. 10.1016/J.CELL.2013.05.039

López-Otín C, Blasco MA, Partridge L, Serrano M, Kroemer G (2023) Hallmarks of aging: An expanding universe. Cell 186:243–278. 10.1016/J.CELL.2022.11.001

Mamun A Al, Shao C, Geng P, Wang S, Xiao J (2024) Recent advances in molecular mechanisms of skin wound healing and its treatments. Front Immunol 15:1395479. 10.3389/FIMMU.2024.1395479

Mascharak S, des Jardins-Park HE, Davitt MF, Griffin M, Borrelli MR, Moore AL, Chen K, Duoto B, Chinta M, Foster DS, Shen AH, Januszyk M, Kwon SH, Wernig G, Wan DC, Lorenz HP, Gurtner GC, Longaker MT (2021) Preventing Engrailed-1 activation in fibroblasts yields wound regeneration without scarring. Science 372:eaba2374. 10.1126/science.aba2374

Moya IM, Halder G (2019) Hippo–YAP/TAZ signalling in organ regeneration and regenerative medicine. Nat Rev Mol Cell Biol 20:211–226. 10.1038/S41580-018-0086-Y

Peña OA, Martin P (2024) Cellular and molecular mechanisms of skin wound healing. Nature Reviews Molecular Cell Biology 25:599–616. 10.1038/s41580-024-00715-1

Porazinski S, Wang H, Asaoka Y, Behrndt M, Miyamoto T, Morita H, Hata S, Sasaki T, Krens SFG, Osada Y, Asaka S, Momoi A, Linton S, Miesfeld JB, Link BA, Senga T, Castillo-Morales A, Urrutia AO, Shimizu N, Nagase H, Matsuura S, Bagby S, Kondoh H, Nishina H, Heisenberg CP, Furutani-Seiki M (2015) YAP is essential for tissue tension to ensure vertebrate 3D body shape. Nature 521:217–221. 10.1038/nature14215

Samaras SE, Almodóvar-García K, Wu N, Yu F, Davidson JM (2015) Global Deletion of Ankrd1 Results in a Wound-Healing Phenotype Associated with Dermal Fibroblast Dysfunction. Am J Pathol 185:96–109. 10.1016/J.AJPATH.2014.09.018

Schlegelmilch K, Mohseni M, Kirak O, Pruszak J, Rodriguez JR, Zhou D, Kreger BT, Vasioukhin V, Avruch J, Brummelkamp TR, Camargo FD (2011) Yap1 acts downstream of α-catenin to control epidermal proliferation. Cell 144:782–795. 10.1016/j.cell.2011.02.031

Shi Y, Reitmaier B, Regenbogen J, Slowey RM, Opalenik SR, Wolf E, Goppelt A, Davidson JM (2005) CARP, a Cardiac Ankyrin Repeat Protein, Is Up-Regulated during Wound Healing and Induces Angiogenesis in Experimental Granulation Tissue. Am J Pathol 166:303–312. 10.1016/S0002-9440(10)62254-7

Tobin DJ (2017) Introduction to skin aging. J Tissue Viability 26:37–46. 10.1016/J.JTV.2016.03.002

Yang JH, Hayano M, Griffin PT, Amorim JA, Bonkowski MS, Apostolides JK, Salfati EL, Blanchette M, Munding EM, Bhakta M, Chew YC, Guo W, Yang X, Maybury-Lewis S, Tian X, Ross JM, Coppotelli G, Meer M V., Rogers-Hammond R, Vera DL, Lu YR, Pippin JW, Creswell ML, Dou Z, Xu C, Mitchell SJ, Das A, O’Connell BL, Thakur S, Kane AE, Su Q, Mohri Y, Nishimura EK, Schaevitz L, Garg N, Balta AM, Rego MA, Gregory-Ksander M, Jakobs TC, Zhong L, Wakimoto H, El Andari J, Grimm D, Mostoslavsky R, Wagers AJ, Tsubota K, Bonasera SJ, Palmeira CM, Seidman JG, Seidman CE, Wolf NS, Kreiling JA, Sedivy JM, Murphy GF, Green RE, Garcia BA, Berger SL, Oberdoerffer P, Shankland SJ, Gladyshev VN, Ksander BR, Pfenning AR, Rajman LA, Sinclair DA (2023) Loss of epigenetic information as a cause of mammalian aging. Cell 186:305–326.e27. 10.1016/J.CELL.2022.12.027

